# Methamphetamine enhances caveolar transport of therapeutic drugs across the rodent blood-brain barrier

**DOI:** 10.1101/2020.05.13.093336

**Authors:** Jui-Hsien Chang, Chris Greene, Clare Futter, Benjamin J. Nichols, Matthew Campbell, Patric Turowski

## Abstract

The blood-brain barrier (BBB) is a multifactorial and multicellular vascular interface separating the systemic environment from the central nervous system (CNS). It gates cerebral penetration of circulating molecules and cells and is the principal reason for low accumulation of many therapeutics in the brain. Low dose methamphetamine (METH) induces fluid phase transcytosis across the BBB *in vitro* and could therefore be used to enhance CNS drug delivery. Here we show, that low dose intravascular METH induced significant leakage exclusively via caveolar transport at the intact BBB in rodents *ex vivo*. Notably, METH-induced leakage was suppressed at 4°C and in Caveolin-1 (CAV1) knockout mice. Furthermore, METH strongly enhanced brain penetration of therapeutic molecules, namely doxorubicin (DOX), a small chemotherapeutic agent, and aflibercept (AFL), a ca. 100 kDA recombinant protein. Lastly, METH improved the therapeutic efficacy of DOX in a mouse model of human glioblastoma (GBM), as measured by a 25% increase in median survival time (p = 0.0024). Collectively, our data indicated that METH can facilitate preclinical assessment of novel experimental treatments and has the potential to enhance drug delivery to the diseased CNS.

## Introduction

The central nervous system is highly vascularised so that the disproportional metabolic demand associated with neuronal activity is fully met (Iadecola, 2017). Whilst meeting the metabolic demands of the CNS, the neuronal vasculature is selectively and dynamically impermeable to protect the delicate ionic neural environment, a feature referred to as the blood-brain barrier. The importance of the BBB is illustrated by its dysfunction in a wide variety of CNS diseases (Sweeney *et al*, 2019).

The BBB is embedded within the neurovascular unit (NVU), comprising vascular endothelial cells, pericytes, astrocytes and neurons, all of which cooperate tightly to establish and regulate the BBB, both during development and its postnatal maintenance (Iadecola, 2017). The physical barrier of the mammalian BBB is mainly provided by the vascular endothelial cells. BBB endothelial cells have tight, virtually impermeable tight junctions sealing the paracellular contacts, and a nearly complete absence of fenestrae and fluid-phase endocytosis (Tietz & Engelhardt, 2015). Thus, large and hydrophilic molecules cannot cross the BBB. To enable entry of nutrients to the CNS, BBB endothelial cells express an array of substrate specific proteins, which either feed into substrate-specific vesicular transport systems or form highly specific channels or membrane transporters (Abbott *et al*, 2010). Small hydrophobic molecules, which may penetrate the NVU, are mostly eliminated by molecular efflux pumps of the ATP-binding cassette (ABC) transporter family of proteins (Saunders *et al*, 2016). Consequently, the BBB constitutes a major impediment for the delivery of therapeutics to the CNS and most drugs do not accumulate at therapeutically required levels in the brain (Abbott, 2013; van Tellingen *et al*, 2015).

Delivery of drugs to the brain has traditionally been investigated in the context of improving chemotherapy-based outcomes for brain tumours, in particular gliomas. Here, the acuteness of the disease allows for a window to establish a relatively close connection between drug transport and therapeutic effect. However, major R & D efforts have also focused on producing efficient transportation of biologicals to treat neurodegenerative diseases, such as Alzheimer’s or Parkinson’s Disease (Goyal *et al*, 2014).

A wide variety of strategies have been explored to enhance drug transport to the brain (van Tellingen *et al*., 2015). Many seek to open the paracellular space between endothelial cells, thus creating a direct passageway between blood and brain parenchyma and enabling enhanced penetration of blood-borne molecules. This can be achieved by osmotically shrinking the endothelial cells (Rapoport, 1988), by focally treating the BBB with a combination of microbubbles and ultrasound (Konofagou *et al*, 2012), by stimulation of the BBB with leakage inducing factors (*e.g*. bradykinin) (van Tellingen *et al*., 2015), and by interfering with endothelial tight junctions or inducing their targeted downregulation (Greene & Campbell, 2016). Undoubtedly, creating a direct connection between brain and the circulation bears significant risks, which are well documented and discussed elsewhere (van Tellingen *et al*., 2015). Therefore, any opening of the BBB to blood constituents needs to be temporally well controlled to avoid significant intoxication of the brain parenchyma with ions and harmful biomolecules.

Alternative BBB drug delivery strategies do not create a direct connection between blood and brain. These include taking advantage of existing receptor-mediated transcytosis (*e.g*. of the Fe-transferrin or insulin receptors) to piggyback antibodies, nanocarriers or AAVs to the brain. Importantly, whilst these strategies leave the BBB physically intact, they require specific adaptation of the drug to transport system that they are targeted towards (Pardridge, 2012). Another important, mainly auxiliary, strategy aims to reduce or block the elimination of the drugs by ABC transporters (van Tellingen *et al*., 2015).

Circulating METH leads to BBB breakdown in rodents (Martins *et al*, 2011; Turowski & Kenny, 2015). Based on this feature, METH has been proposed for use to enhance drug transport to the diseased brain (Kast, 2007). METH induces BBB breakdown in an endothelial cell autonomous or non-autonomous fashion, each governed by a distinct mechanism of BBB opening: At concentrations above 10 μM, METH leads to - often chronic-endothelial junction breakdown with slow onset (Coelho-Santos *et al*, 2015). This likely occurs in response to inflammatory signalling in the endothelium, triggered either by METH acting directly on the EC or due to its cytotoxic effect on the neuroglial environment. In contrast, METH at concentrations in the low micromolar range leaves endothelial junctions intact and instead rapidly induces fluid-phase transcytosis in cultured BBB endothelial cells (Martins *et al*, 2013).

Since METH at low concentrations does not induce paracellular opening we sought to investigate this mechanism further and corroborate its relevance at an intact NVU. We used perfused rodent brains *ex vivo* to show that fluid-phase transcytosis occurred in the intact brain in response to METH and that this involved transport-competent caveolae. We also investigated the proposed utility of METH as a facilitator for BBB drug transport and showed for the first time that METH enhanced transport of DOX and the anti-VEGF trap aflibercept. Lastly, the efficacy of the common chemotherapeutic drug, DOX, to treat human GBM bearing mice was enhanced by co-treatment with METH.

## Results and Discussion

We developed an *ex vivo*, dual carotid perfused model of the intact rat brain, to study BBB leakage in highly controlled fashion (Supplemental Figure 1A). When rat heads were perfused via both common carotid arteries at equal pressure, perfusate constituents were durably kept within the side of the head, which they were applied to, indicating that under these experimental conditions mixing did not occur at the Circle of Willis (Supplemental Figure 1B). Thus, this model allowed to study an experimental condition and its control within the same brain. Preparation of rodent heads and the perfusion protocol was identical to that used to study a fully functional BRB in explants of retinae (Warboys *et al*, 2009), suggesting that this protocol left the BBB intact. Indeed, when heads were perfused with Evans Blue/albumin (EB/Alb), this dye remained restricted to the vasculature of most areas of the brain, including cortex, hippocampus and thalamus, and did not leak for at least 1 h (Supplemental Figure 1C), indicating that BBB properties were preserved.

To analyse leakage across the BBB, METH (1 μM) was included with the EB/Alb-containing perfusate in one side of *ex vivo* brains. After 60 min, the entire head vasculature was cleared to remove intravascular dye. Subsequent analysis of fixed brain slices revealed strong and widespread accumulation of EB/Alb but only in the in the hemisphere METH was administered to (Figure 1 A, B; Supplemental Figure 2A). The accumulated EB/Alb was located outside of the vasculature (Supplemental Figure 2 B, C), indicating that leakage and BBB breakdown had occurred. METH-induced leakage was completely suppressed at 4°C (Figure 1 C, Supplemental Figure 2 D, E), demonstrating that it was cold-sensitive and thus likely to be dependent on vesicular traffic (Boulenc *et al*, 1993; Jungmann *et al*, 2008). In cultured BBB endothelial cells METH induces permeability via small pinocytic vesicles with diameters of 70 to 100 nm, reminiscent of caveolae (Martins *et al*., 2013). Double carotid perfused brains were treated with or without METH and HRP as leakage tracer and brain microvessels analysed by DAB-EM (Figure 1 D,E). In METH-treated sides the immediate environment of microvessels displayed perivascular oedema with notable astrocyte endfoot swelling. Accordingly, vessels appeared compressed, with a less rounded appearance and reduced diameter. Paracellular junctions between endothelial cells appeared ultrastructurally identical to those found in the contralateral control side and did not display any accumulation of DAB, indicating that tight junctions were left intact. However, microvascular endothelial cells in METH-treated hemispheres contained large number of DAB positive vesicles < 100 nm diameter, which were uncoated and thus resembled caveolae. Quantification revealed that endothelial accumulation of these vesicles was highly specific to the METH treated side (Figure 1 F). In the METH-treated side, the number of DAB-positive endothelial vesicles < 100 nm was significantly induced. The number of vesicles > 100 nm was also significantly increased but much less strongly. Next, we sought direct proof of caveolae being responsible for BBB tracer leakage in response to METH in our *ex vivo* brain model. For this, the double carotid perfusion model was adapted for use in mice. In wild type mice METH induced leakage, which was similar to that seen in rats (Figure 1 G; Supplemental Figure 3). In contrast, in Cav1 k/o mice, which lack caveolae (Razani *et al*, 2001), there was a complete absence of METH-induced leakage. Thus, we concluded that as in cultured BBB endothelial cells, METH induced leakage at the intact NVU via transport competent caveolae. This was in agreement with other recent studies demonstrating the initial phase of BBB leakage during experimental stroke is mediated by caveolae (Knowland *et al*, 2014). Furthermore, lack of pericytes or the pericyte-induced endothelial lipid transporter MFSD2A leads to upregulation of caveolae at the BBB and leakage (Andreone *et al*, 2017). Lastly, transcellular lymphocyte migration across the BBB also requires caveolar transport (Lutz *et al*, 2017).

**Figure 1.**
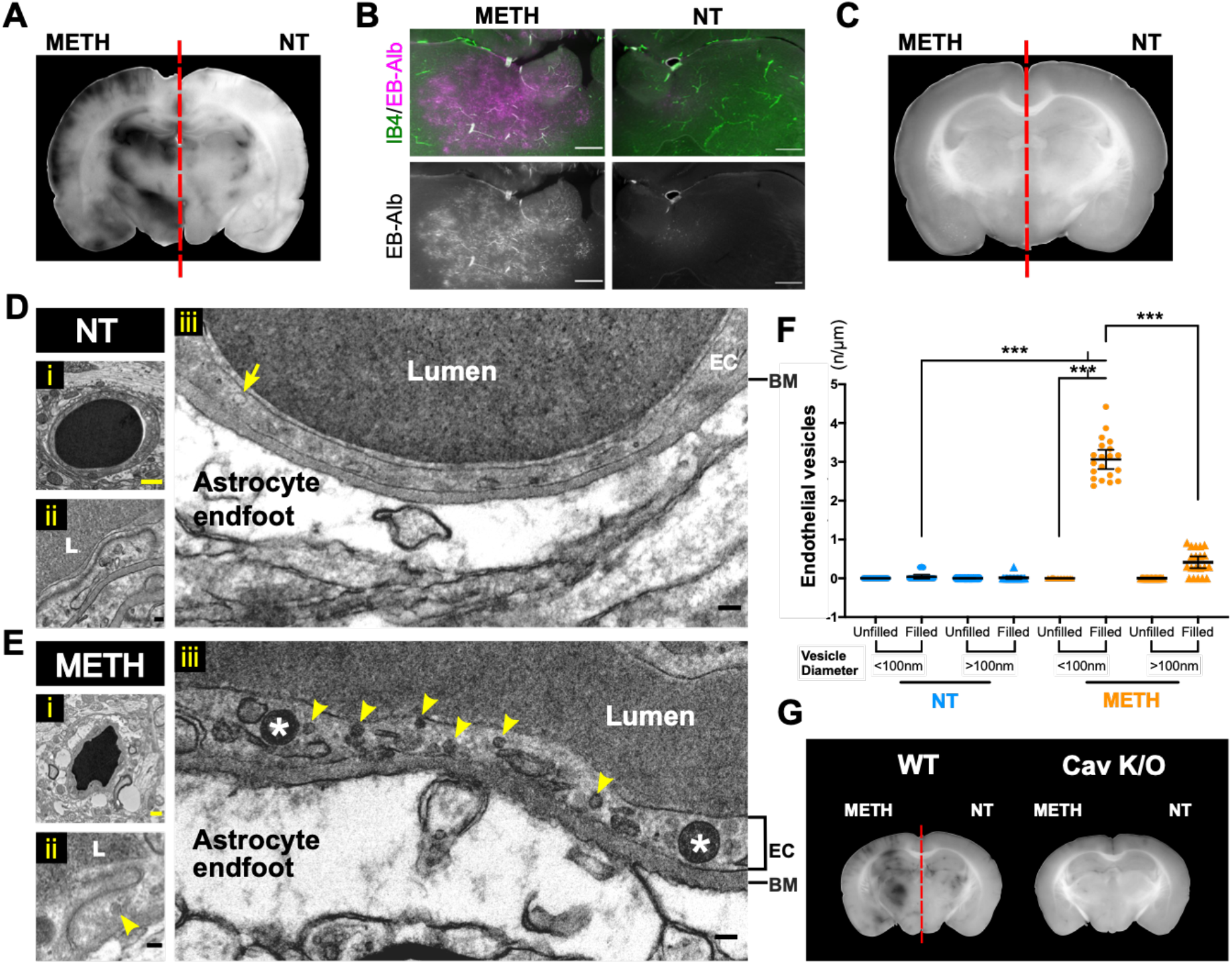
METH induces BBB transport via endothelial caveolae. (**A, B**) Rat brains were perfused *ex vivo* through both carotid arteries with EB/Alb as illustrated in Supplemental Figure 1. METH (1 μM) was included in the perfusate in the left carotid artery. After 60 min EB/Alb was flushed from the vasculature and brains perfused fixed, sectioned and analysed. Shown in (A) is a brightfield image of a representative coronal section with a full sectioning profile provided in Supplemental Figure 2A. Note large parenchymal areas with accumulated EB in the METH but not NT side of the brain (except in the lateral ventricular areas). Shown in (B) are corresponding fields from the upper medial thalamus, imaged by fluorescent microscopy with EB in magenta and the vasculature (counterstaining with IB4) in green-FITC. Scale bars, 500 μm. (**C**) As in (A) except that head was precooled to 4°C and perfused with ice-cold solutions. A full sectioning profile is shown in Supplemental Figure 2D. (**D, E, F**) As in (A) but using HRP as a leakage tracer. HRP was not flushed from the vasculature before fixation. Subsequently, corresponding areas in brain sections were analysed by DAB-EM. Shown in (D, E) are vertical sections through representative HRP microvessels in the thalamus and cortex from NT (D) or METH-treated sides (E). Panels (i) represent overviews of the vessels, for which magnified intact endothelial paracellular junction areas are shown in (ii) (L = lumen), and ca. 3 μm vascular sections in (iii). Arrow: empty intraendothelial vesicle < 100 nm; Arrowheads: HRP-filled intraendothelial vesicles < 100 nm; Stars: HRP-filled intraendothelial vesicles > 100 nm; EC: endothelial cell; BM: basement membrane. Note the marked enlargement of astrocyte endfeet in response to METH treatment (E), indicating severe perivascular oedema. (F) Vesicle count per μm of endothelial plasma membrane length determined from EM images as in (D, E). Data from three independent brains and a total of 20 microvessels. ***, p<0.001 (ANOVA). Yellow scale bar: 1 μm; Black scale bar: 1 nm. (**G**) As in (A) but performed in either wt or Cav k/o mice. Note the complete absence of METH-induced leakage in Cav k/o brains.

We found METH-induced leakage to EB/Alb and HRP to be very similar. We next explored the possibility that METH could enhance BBB transport of therapeutic molecules with different physical and chemical properties. For this, we used DOX and the anti-VEGF trap aflibercept as paradigms for small chemotherapeutic molecules and large therapeutic proteins, respectively. Both were applied to rodent heads via the dual carotid method in the presence and absence of 1 μM METH. After 60 min, brains were isolated and examined for the presence of either therapeutic molecule. In the absence of METH, DOX autofluorescence was virtually absent from the brain parenchyma (Figure 2A). In contrast, in the presence of METH, strong DOX autofluorescence was detected on capillaries and throughout the entire brain. Similarly, aflibercept, as detected by indirect immunochemical staining, was mainly found associated with vascular lumens and very few occurrences of focal parenchymal accumulation in the absence of METH. In contrast, most areas of the METH-treated hemispheres showed a dramatic increase of diffuse parenchymal aflibercept accumulation. Collectively, these results showed that DOX and aflibercept were excluded from the brain by a functional BBB and that METH enhanced their transport to the brain, suggesting that the BBB delivery of a wide variety of circulating molecules could be enhanced by METH.

**Figure 2.**
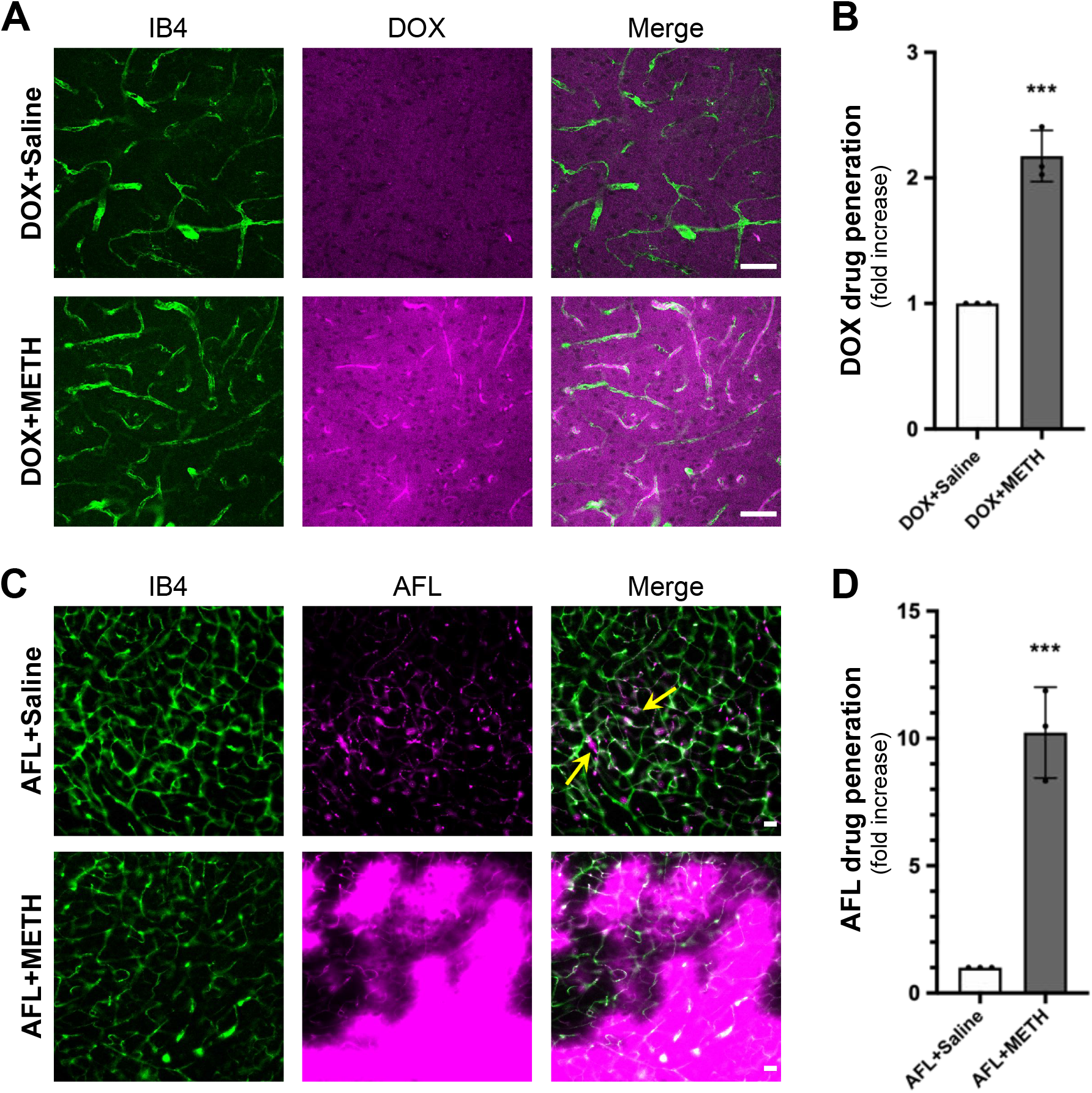
METH enhances drug transport across the BBB. (**A, C**) Rat brains were perfused *ex vivo* through both carotid arteries with cardioplegic solution containing DOX (10 μg/mL)(A) or AFL (0.5 mg/mL) (B) and METH (1 μM) v saline in opposing sides as indicated. After 60 min the vasculature was flushed and brains perfused fixed, sectioned, stained, analysed and quantified. Shown are representative fields from thalamus stained using IB4 (green). DOX was detected by virtue of its autofluorescence (A, magenta), whilst AFL was revealed by staining using goat polyclonal anti-human IgG Fc antibody (B, magenta). Scale bars, 50 μm. (**B, D**) Quantification of drug penetration. Data from 3 individual brain sections of DOX- or AFL- treated vasculature with its corresponding control. ***, p<0.001 (t-test).

Next we studied if METH enhanced the therapeutic efficacy of DOX in a mouse model of human glioblastoma (GBM). Previous studies suggest that caveolae-mediated BBB opening is only seen following exposure of BBB endothelial cells to METH in the low micromolar range. We carried out a basic pharmacokinetic study in mice to determine how i.p. METH injection translated to METH accumulation in plasma over time. Plasma METH concentrations peaked ca. 30 min following i.p. injection, declining to nearly 50 % after 60 min (Supplemental Figure 4), in line with a previous report examining the pharmacokinetics of METH following i.p. administration at 30 mg/kg ((Martins *et al*., 2011). Modelling suggested that METH injected i.p. at 2.5 mg/kg led to METH at plasma concentrations of ca. 1 μM for at least 1 h.

Brains of Balb/c nude mice were injected with human D270 cells to induce robust formation of tumours bearing many features of human GBM (Miao *et al*, 2014). Treated with a standard regimen of DOX, such mice survive for ca. 18-19 days. D270-inoculated Balb/c mice were treated with either DOX (6mg/kg) or DOX and METH (2.5 mg/kg) on days 3, 6, 9, 12 and 15 after tumour cell inoculation. A significant difference in survival of the mice undergoing the two different treatment regimens was observed (Figure 3 A). Median survival in the DOX only group (n=17) was 18 (± 1.1) days and in line with previous reports for this model (Miao *et al*., 2014). In contrast, median survival time in the DOX+METH group (n=20) was 22.5 (± 1.5) days, i.e. significantly increased by ca. 25 % (p = 0.0024).

**Figure 3.**
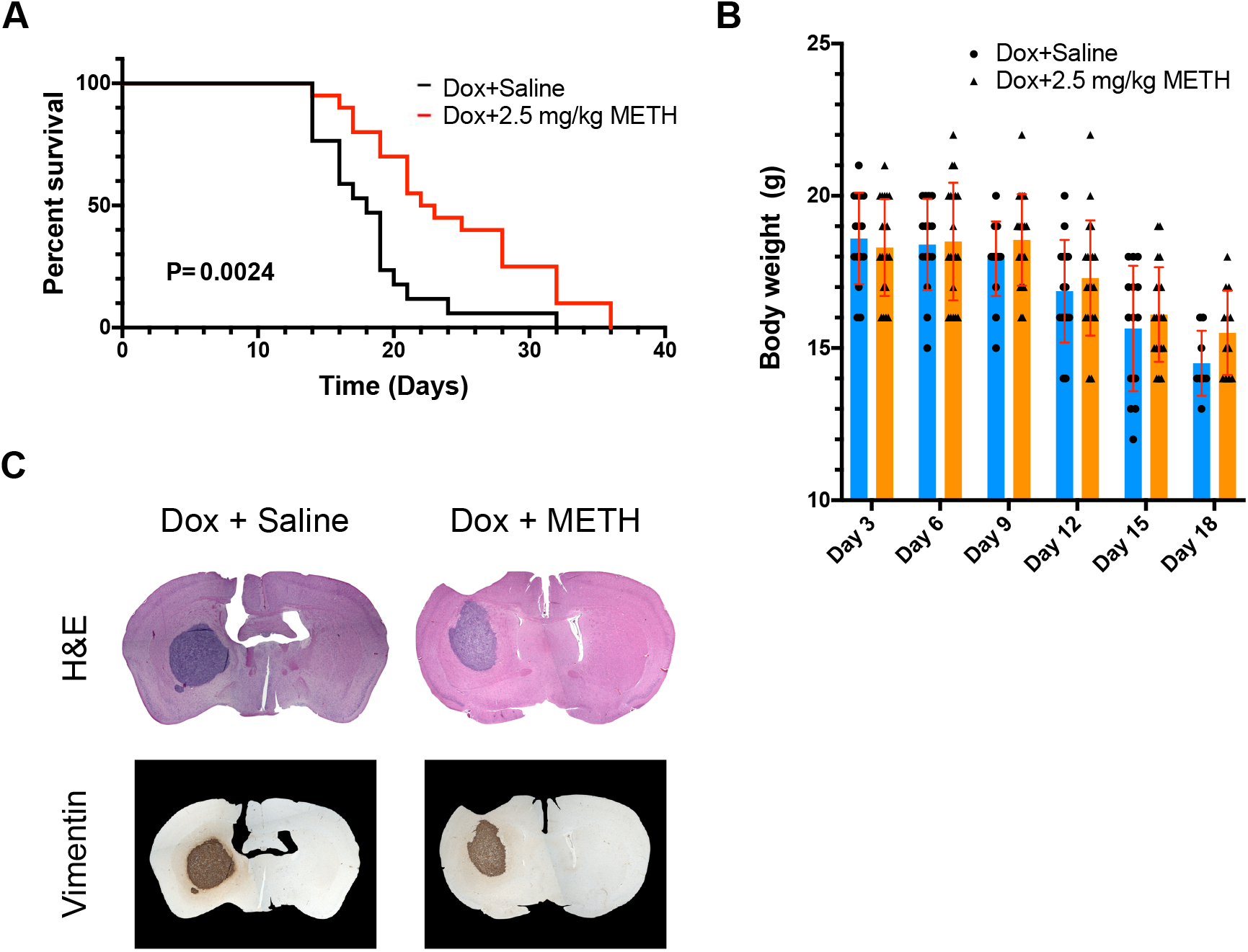
METH enhances Dox-mediated survival in glioma bearing mice. (**A, B**) The right caudate-putamen region of athymic nude mice were injected with human D270 cells. Subsequently, all mice were treated i.p. with 6 mg/kg Dox on days 3, 6, 9, 12 and 15. Additionally, mice were randomised and received an additional i.p. injection of saline or METH (2.5 mg/kg). Survival was recorded and is shown (A). Body weight was also recorded for both experimental groups and is shown as mean ± SD in (B). (**C**) Representative coronal sections of brains, removed and stained from animals at time of death as described for panel (A). Brains were either stained using H & E, or against human vimentin and showed similar expansion of the tumour at time of death.

Importantly, weight loss in both groups was indistinguishable, indicating that differences in survival times were not due to METH improving general health independently (Figure 3 B). This survival improvement was similar to that seen when D270 tumour grafts are targeted with specifically designed chimeric antigen receptor T cells (Miao *et al*., 2014). Post mortem histological analyses of brain sections showed the presence of very large tumour growths, which were clearly identifiable by H & E staining or staining using anti-human vimentin antibodies (Figure 3C). Tumours were similar in size and appearance in both, the DOX only and DOX+METH groups.

In conclusion, in this proof-of-principle study we have demonstrated that METH triggered caveolar transport of proteins and small molecules at the intact neurovascular unit and that this process could be harnessed to enhance brain availability of a circulating drug. Therapeutic use of METH can be safe and its use is FDA approved to treat obesity as well as Attention Deficit Disorder with Hyperactivity (https://www.accessdata.fda.gov/drugsatfda_docs/label/2013/0005378s027lbl.pdf). Our study showed that induction of BBB opening occurred with METH at low concentrations and thus compatible with relative safe use. Other reported METH treatment regimens employ much higher concentrations, which are demonstrably more neurotoxic in rodents and likely to be so in humans (also discussed in Turowski & Kenny, 2015). Furthermore, induction of transport-competent caveolae at the BBB offers clear advantages over opening paracellular junctions, the mechanism invoked of treatment regimens utilising higher METH concentrations. It allows transport of blood constituents to the brain parenchyma without creating a direct connection between blood and brain, and thus displays much lower toxicity for neuronal networks and in particular their delicate ionic environment. Lastly and most importantly, we showed, for the first time, clear therapeutic benefits for METH use in combination with a chemotherapeutic drug in a mouse glioma model, as measured by survival times. Residence time of METH in the circulation was very short-lived in mice (Martins *et al*., 2011) (see also Supplemental Figure 4) as it is in humans (Kuypers *et al*, 2016), pointing towards a short window of opportunity, during which the BBB was opened for blood-borne molecules. Thus, BBB opening by METH constitutes a temporally well controlled process, highlighting a highly desirable feature due to potentially reduced brain toxicity. We propose that adjunctive METH treatment could be rapidly developed for versatile clinical use. Indeed, our data suggested that METH-induced caveolae will transport molecules of widely varying chemical composition and size, raising the likely possibility that even transport of nucleic acids and viral particles to the brain can be enhanced by low level METH. Use of METH as an adjuvant will facilitate preclinical assessment of novel experimental treatments. Furthermore, in conjunction with chemotherapy it may be of benefit to GBM patients, but also have value for other disease scenarios and treatment modalities, where the BBB continues to be a significant therapeutic impediment, such as rare paediatric cerebral and many neurodegenerative diseases.

## Material and Methods

### Animals

Adult female Wistar rats and C57BL6 mice were purchased from Charles River Laboratories Inc. (Oxford, UK). Tie2-GFP Cav-1 K/O mice were provided by Dr. Ben Nichols (LMB, Cambridge). In Caveolin 1-/-mice, CAV1 gene was disrupted by a designed targeting vector replacing the first 2 exons by a neomycin resistance cassette (Razani *et al*., 2001). All experiments using animals were conducted in accordance with the UK Home Office Guide on the Operation of Animal (Scientific Procedures) Act of 1986 with appropriate licenses.

### *In situ* dual carotid perfusion assay

After CO_2_ asphyxiation, external and internal jugular veins of rats or mice were transected bilaterally. Both common carotid arteries were isolated and cannulated using polyurethane cannulae (3.5 Fr for rat and 1 Fr for mouse). The head vasculature was immediately flushed with heparin (300 U/mL in saline) and then with a cardioplegic solution (10 mM MgCl_2_, 110 mM NaCl, 8 mM KCl, 10 mM HEPES, 1 mM CaCl_2_ and 10 μM isoproterenol, pH 7) in order to reduce thromboembolism, preserve tissue viability, and stabilise the head vasculature. Subsequently, heads were perfused via bilateral common carotid cannulae with cardioplegic solution containing the leakage tracers, EB/Alb (EB-Alb; 5 mg/mL Evans Blue, 10% albumin) or horseradish peroxidase (HRP; 5 mg/mL), and optionally IB4, METH, DOX or AFL. METH was given at 1 μM. After incubation of treatments, the remaining intravascular perfusate was flushed out and the vasculature was cleaned by perfusion of cardioplegic solution. Before removal of brain and eyes, perfusion fixation was carried out with 4% Paraformaldehyde (PFA) or Karnovsky EM fixatives (3% v/v glutaraldehyde, 1% v/v PFA in 0.08M sodium cacodylate buffered to pH 7.4 with 0.1M HCl) for IHC or EM studies, respectively.

All solutions were administrated simultaneously to both hemispheres with an equal delivery pressure throughout the experiments.

### Immunohistochemistry and histology

#### Ex vivo brains and retinae

After PFA perfusion fixation, *ex vivo* brains from *in situ* dual perfusion assay were immersed in 4% PFA for 24 h, sectioned into 100 μm and 500 μm slices by Vibratome^®^ 1000 Plus Sectioning System (FEDELCO, S.L.). Retinae were isolated from the eyes and fixed in PFA for 1 h before further processing. 100 μm brain sections and retinae were blocked with 1% FBS/2X PBS (3% Triton x-100, 0.5% Tween 20, 0.2% sodium azide) and stained overnight at 4C with FITC-conjugated IB4 (Fluorescein labeled Griffonia Simplicifolia Lectin I isolectin B4, FL-1201, Vector laboratories, 1:200) or primary antibody against Cld-5 (mouse monoclonal, C43C2, 35-2500, Thermo Fisher scientific, 1:200). AFL was labeled at the Fc portion of human IgG immunoglobulin (goat polyclonal anti-human IgG Fc antibody, NB7446, Novus Biologicals, 1:200 at 4C overnight). Stainings were followed by a corresponding Alexa Fluor-conjugated secondary antibody (Thermo Fisher Scientific, 1:200) at room temperature for 2 h.

#### GBM xenograft brains

Whole brains previously cryo-fixed using OTC were immersed in 10% neutral buffered formalin and embedded in paraffin. Following preparation of the paraffin sections, samples were dewaxed, rehydrated, stained with Harris Haematoxylin (Pioneer Research Chemicals) and 1% Eosin (Pioneer Research Chemicals) by an automated slide stainer. Alternatively, human vimentin immunohistochemical staining of the paraffin sections was performed by using the Ventana Discovery XT instrument (Roche) with Ventana DAB Map detection Kit (760-124, Roche) and pre-treatment of an EDTA equivalent, cell conditioning 1 (CC1, 950-124, Roche). Glioma graft cells were labeled by the anti-human Vimentin antibody (mouse monoclonal, V9, 790-2917, Roche, pre-diluted), followed by a mouse secondary antibody (rabbit polyclonal anti-mouse biotin antibody, E0413, Dako, 1:200). Histopathological studies of GBM xenograft mice were conducted by Division of Neuropathology, UCL Queen Square, Institute of Neurology.

### Transmission electron microscopy (TEM)

After perfusion fixation with Karnovsky solution, *ex vivo* brains treated with HRP-containing perfusate (5 mg/mL) were immersed in the same EM fixative for no more than 48 h, sectioned into 100 μm slices by Vibratome^®^. The 100 μm free-floating sections were incubated in diaminobenzidine (DAB) solution (0.075% DAB/0.02% hydrogen peroxide in 0.1M Tris) for 30 mins at room temperature in the dark. After the DAB reaction, specimens were secondarily fixed in 1% Osmium tetroxide/1.5% Potassium ferrocyanide (in H2O) for 1.5 h at 4C in dark. They were further dehydrated by EtOH, infiltrated with propylene oxide (PO) and finally embedded in Epoxy resin after polymerization of Epon overnight at 60C. Sample blocks were cut into 70 nm ultrasections by Leica EM UC7 Ultramicrotome, mounted onto 300 mesh EM grids and stained with lead citrate.

### Imaging

Gross images of 100 and 500 μm brain sections were acquired by a stereo fluorescence microscopes (Olympus) equipped with a colour camera. H&E and Vimentin stained sections were scanned by EVOS FL auto 2 imaging system (Thermo Fisher Scientific). High magnification fluorescent microscopy was performed either on an Axioscope (Carl Zeiss), or by confocal laser scanning microscopy on a CLSM 700 (Carl Zeiss). JEM-1400 Transmission Electron Microscope (JOEL) equipped with a digital camera (Gatan Orius) was used for all TEM studies. Images were processed by ImageJ (NIH).

### TEM quantification

Total 3 *ex vivo* rat brains underwent EM analysis for quantification of vesicular density. Vessels < 10 μm in diameter were imaged from the METH-treated hemispheres and their corresponding control hemispheres. HRP-filled and non-HRP-filled vesicles were counted per EC length (n/μm) and categorized into 4 groups: > 100 nm filled vesicles, < 100 nm filled vesicles, > 100 nm filled vesicles and < 100 nm un-filled vesicles, according to the filling of tracers and its diameter.

### METH dosing *in vivo*

Total 15 C57BL6 mice, weighting 17.5 g in average, were randomly allocated to 5 dosing groups (3 mice per group): 0.75 mg/kg, 2.5 mg/kg and 7.5 mg/kg, treated for 60 min by peritoneal injection; and an additional 7.5 mg/kg group with a shorter treatment time of 30 min. Plasma METH concentrations was measured by LC-MS/MS (conducted by Cyprotex). A pharmacokinetic approximation was achieved by fitting the data to a quadratic function. Based on the hypothetical pharmacokinetic curve, peak plasma concentrations of METH were determined.

### Stereotaxic injection of D270 cells into athymic nude mice

D270 cells were isolated from a glioblastoma multiforme tumour in Duke University as previously described (Humphrey *et al*, 1988). D270 cells were cultured in improved MEM zinc option media containing 2.383 g/L HEPES buffer, 5 mg/L insulin, 584 mg/L L-glutamine, 5 μg/L selenium and 2.2 g/L sodium bicarbonate. Balb/c nude mice (CAnN.Cg-Foxn1nu/Crl) were anaesthetised with a mixture of medetomidine/ketamine and placed in a stereotaxic frame. A 1 cm midline incision was made in the scalp and a burr hole was drilled above the frontal cortex (coordinates from bregma: A/P = +0.5 mm; M/L = −2.5 mm; D/V = +3.0 mm). A Hamilton syringe containing 3 μl of cell suspension (1×10^5^ cells) was slowly lowered into the brain and cells were injected at a rate of 0.2 μl/min. The needle was left in place for 5 min to prevent reflux. Animals were sutured and placed in an incubator until they had recovered. All mice received an intraperitoneal injection of DOX (6 mg/kg in saline) on days 3, 6, 9, 12 and 15. Prior to doxorubicin injection, animals were randomised and received an intraperitoneal injection of saline or methamphetamine (2.5 mg/kg in saline). Mice were sacrificed and brains were removed and fixed in 4 % formaldehyde overnight at 4°C, washed 3x in PBS and cryoprotected in 10 %, 20 % and 30 % sucrose before being snap-frozen in OCT compound.

### Statistics

Statistical analyses were carried out by Microsoft excel (Office 365) or Prism 8 (GraphPad). Experiment sample sizes were determined empirically. Two and multiple group comparisons were analyzed by unpaired two-tailed Student’s t test and one-way ANOVA, respectively. Survival data were analysed by using a Mantel-Cox Log-rank test.

## Acknowledgements

This work was supported by grants from the British Heart Foundation (FS/16/26/32193) and Cancer Research UK (C26070/A24762).

## Author contributions

J-H.C.: Conceptualization; Data curation; Formal Analysis; Investigation; Methodology; Visualization; Writing – original draft

C.G.: Data curation; Formal Analysis; Investigation; Methodology; Visualization

C.F.: Conceptualization; Data curation; Formal Analysis; Investigation; Methodology

B.J.N.: Conceptualization; Resources

M.C.: Conceptualization; Resources Data curation; Formal Analysis; Methodology; Supervision; Writing – original draft

P.T.: Conceptualization; Data curation; Formal Analysis; Project administration; Funding acquisition; Resources; Supervision; Writing – original draft

## Conflict of interest

The authors declare no conflict of interest.

## Supplemental Figures

**Supplemental Figure 1.**
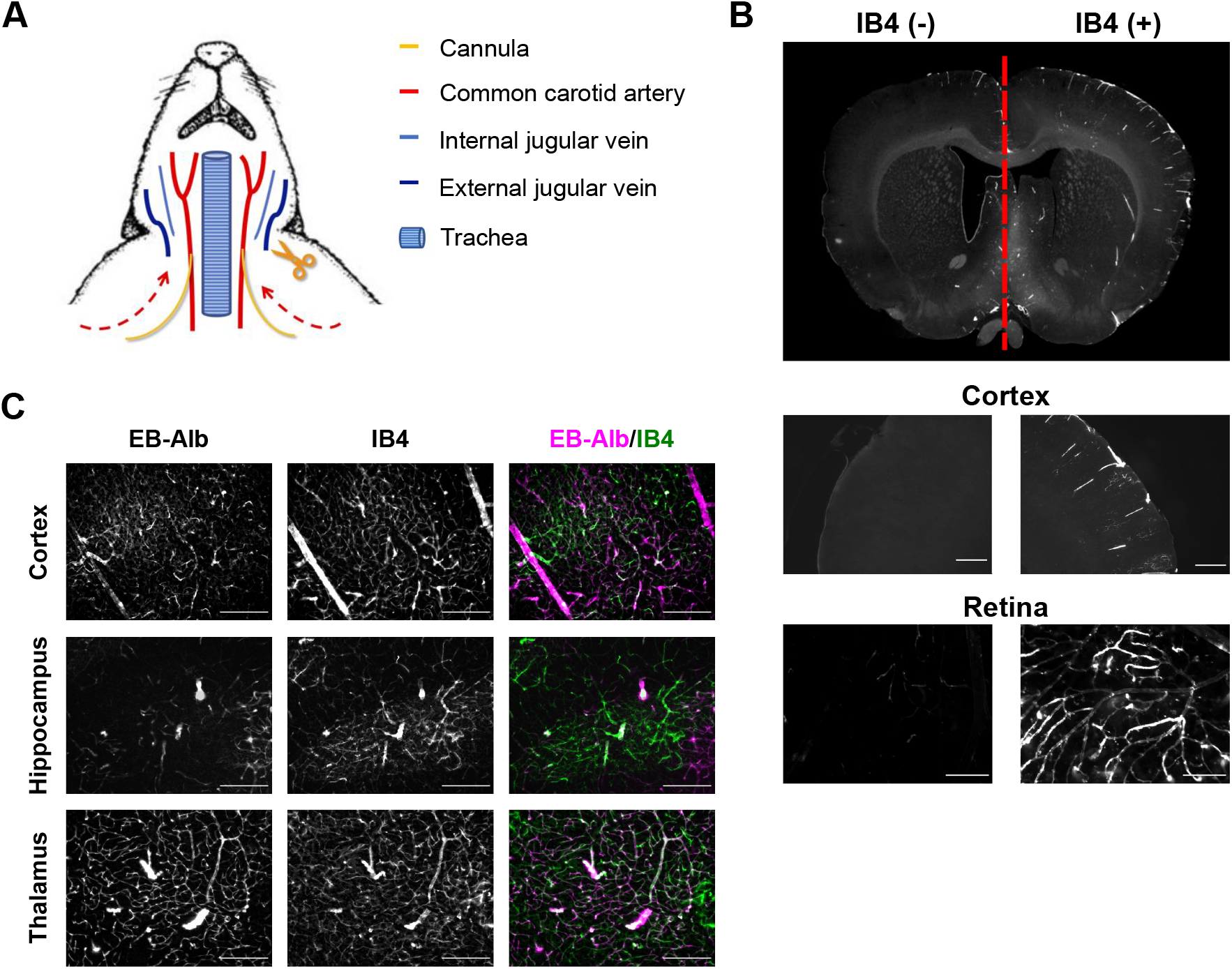
The dual carotid artery perfusion model. (**A**) Schematic of the experimental setup for rats and mice. Shortly after death, the two common carotid arteries were cannulated and the jugular veins sectioned. Heads were then perfused with saline containing heparin, followed by a cardioplegic solution, shown to preserve the vasculature and the BBB (Warboys *et al*., 2009). Subsequently, heads were incubated by perfusion with cardioplegic solution containing tracer molecules, METH, DOX, and aflibercept (or in control sides their respective vehicles). (**B**) Both sides of rat heads were perfused at equal pressure. DyLight^®^ 594 GSL IB4 was included in the perfusate of the right carotid artery. After 1 h, heads were perfused fixed, brains isolated and sections examined by epifluorescence microscopy. Note that IB4 only appeared in the right brain hemispheres and the right retinal vasculature, demonstrating that mixing of perfusates did not occur at the Circle of Willis. (**C**) Rat heads were perfused using the dual carotid artery model as described. EB/Alb was included with the perfusate and left within the heads for 1 h. Heads were perfused fixed, sectioned and the vasculature stained using IB4 before analysis by epifluorescent light microscopy. Note that EB was retained within the vasculature in all brain areas shown, indicating preservation of the BBB.

**Supplemental Figure 2.**
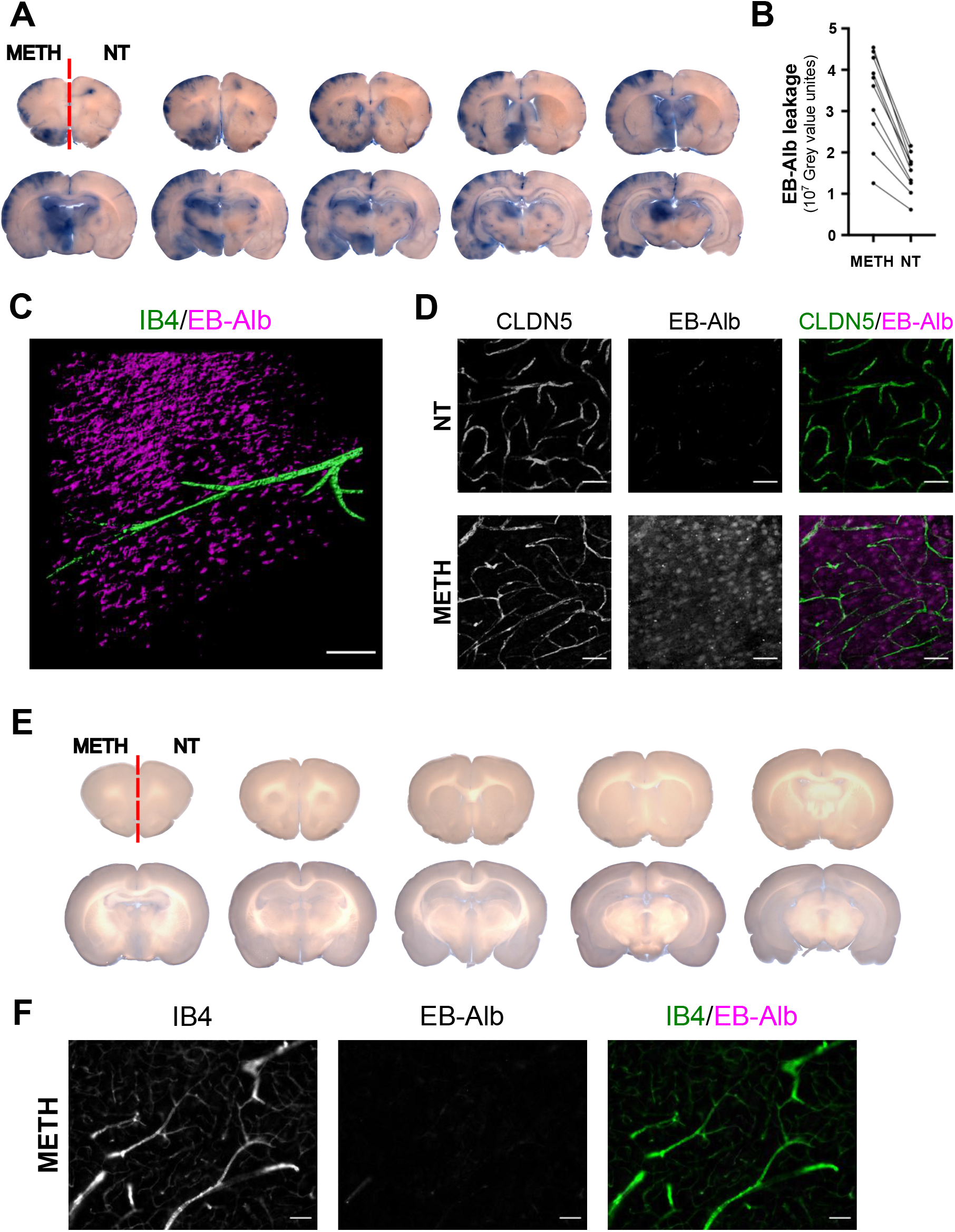
METH-induced leakage in the dual carotid artery perfused rats. (**A, B**) As Figure 1 A, (A) showing the full set of coronal sections and the EB-Alb leakage of the METH- and non-treated vasculatures from each individual section was quantified in (B). (**C**) As Figure 1 B, showing a 3-D rendering of confocal stacks from a representative branching microvessel and demonstrating extravasation of EB/Alb into the parenchyma. Scale Bar, 10 μm. (**D**) As in Figure 1 B but counterstained for CLDN5, indicating integrity of CLDN5 and extravasation of EB/Alb. Scale Bar, 50 μm. (**E**) As Figure 1 C, showing the full set of coronal sections. (**F**) Brains were treated cold as described for Figure 1 C and coronal sections counterstained using IB4 (green). Shown are METH-treated fields from thalamus. Scale bars, 10 μm.

**Supplemental Figure 3.**
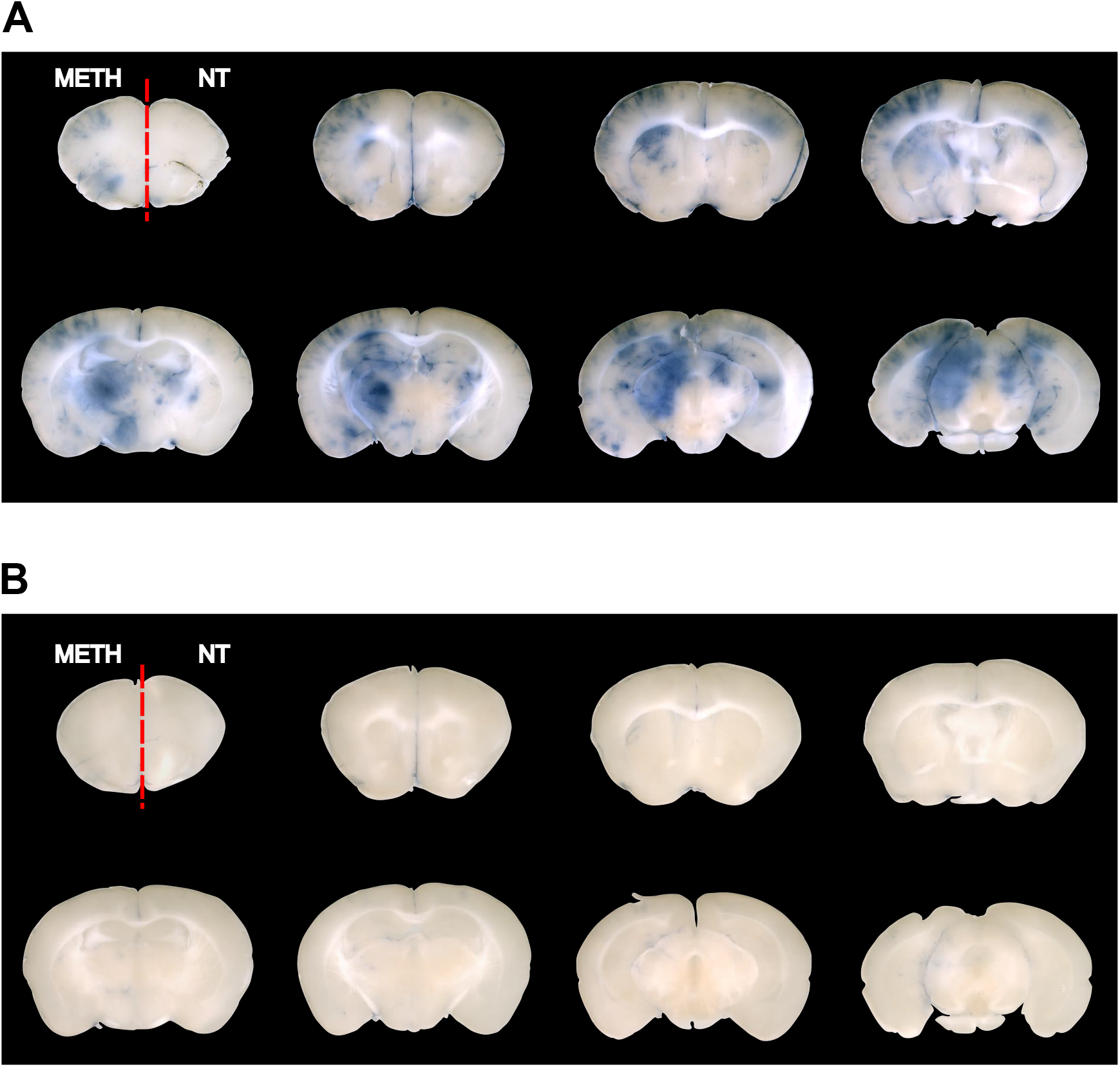
METH-induced leakage requires Caveolin 1. (**A**) As Figure 1 G, showing a full set of representative coronal sections from wt mice. (**B**) As Figure 1 G, showing a full set of representative coronal sections from Cav k/o mice.

**Supplemental Figure 4.**
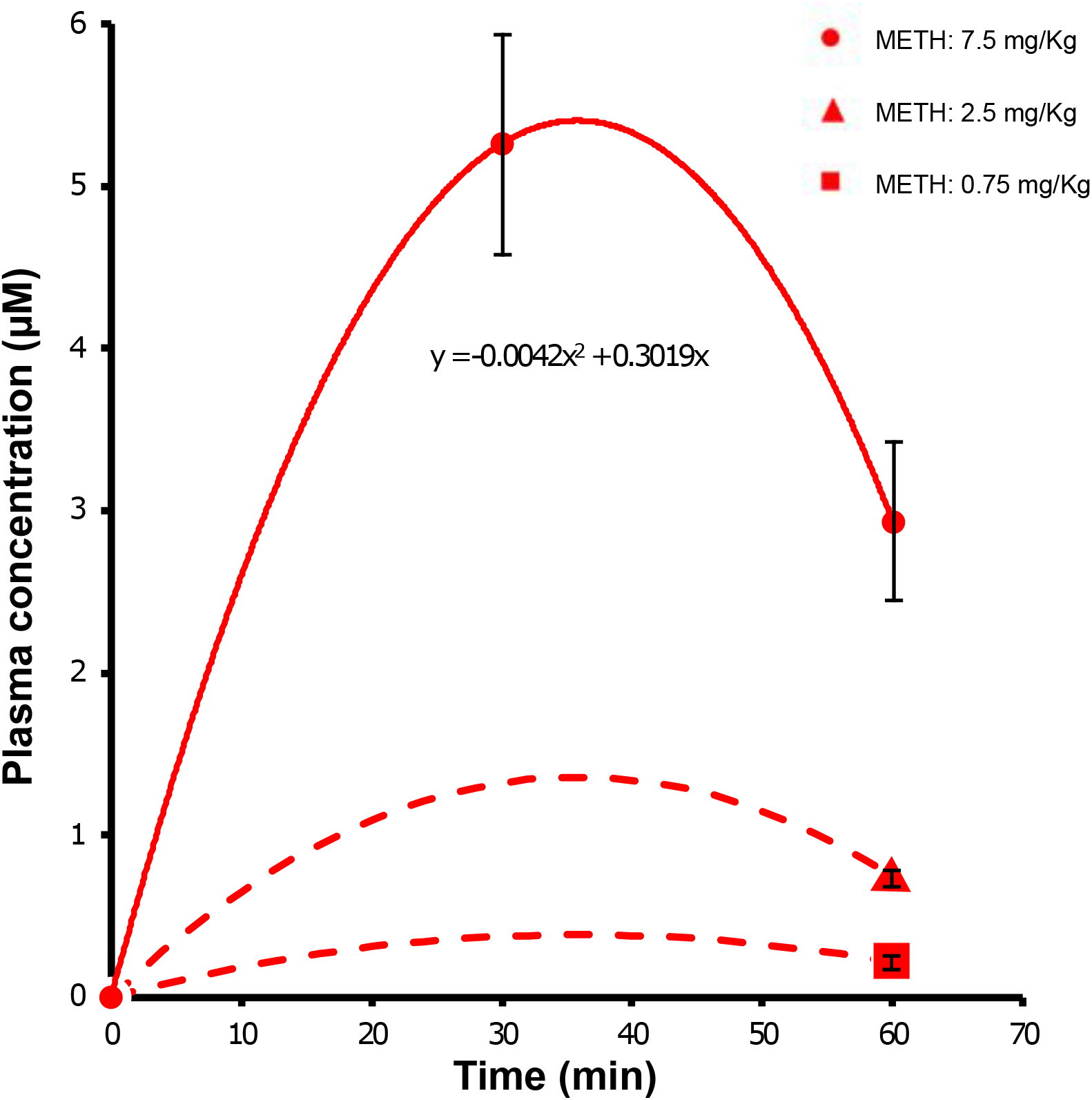
Estimation of pharmacokinetics of METH in mice. C57BL6 mice were injected i.p with the indicated doses of METH. Plasma was isolated at either 0, 30 or 60 min and METH content determined by LC-MS/MS. Data points are means ± SEM from 3 independent experiments. Values from the 7.5 mg/kg data set was then fitted to the polynomial function y=a*x^2^+b*x, resulting in the fitting parameters a = 0.0042, b = 0.3019 (R^2^ = 1). The time for maximal plasma concentration was determined from the derivation y = 2*a*x + b = 0 as 35.9 min, which was in good agreement with an earlier report (Martins *et al*., 2011). Curve estimation for plasma METH concentration following injections of 2.5 and 0.75 mg/kg were generated by using the experimental data points at 0 and 60 min and assuming that the peak occurred at the mathematically time of 35.9 min. Additional data points were not taken to comply with the principles of the 3Rs (http://www.3rs-reduction.co.uk).

## Notes

### Competing Interest Statement

The authors have declared no competing interest.

